# HiC-bench: comprehensive and reproducible Hi-C data analysis designed for parameter exploration and benchmarking

**DOI:** 10.1101/084954

**Authors:** Charalampos Lazaris, Stephen Kelly, Panagiotis Ntziachristos, Iannis Aifantis, Aristotelis Tsirigos

## Abstract

**Background:** Chromatin conformation capture techniques have evolved rapidly over the last few years and have provided new insights into genome organization at an unprecedented resolution. Analysis of Hi-C data is complex and computationally intensive involving multiple tasks and requiring robust quality assessment. This has led to the development of several tools and methods for processing Hi-C data. However, most of the existing tools do not cover all aspects of the analysis and only offer few quality assessment options. Additionally, availability of a multitude of tools makes scientists wonder how these tools and associated parameters can be optimally used, and how potential discrepancies can be interpreted and resolved. Most importantly, investigators need to be ensured that slight changes in parameters and/or methods do not affect the conclusions of their studies.

**Results:** To address these issues (compare, explore and reproduce), we introduce HiC-bench, a configurable computational platform for comprehensive and reproducible analysis of Hi-C sequencing data. HiC-bench performs all common Hi-C analysis tasks, such as alignment, filtering, contact matrix generation and normalization, identification of topological domains, scoring and annotation of specific interactions using both published tools and our own. We have also embedded various tasks that perform quality assessment and visualization. HiC-bench is implemented as a data flow platform with an emphasis on analysis reproducibility. Additionally, the user can readily perform parameter exploration and comparison of different tools in a combinatorial manner that takes into account all desired parameter settings in each pipeline task. This unique feature facilitates the design and execution of complex benchmark studies that may involve combinations of multiple tool/parameter choices in each step of the analysis. To demonstrate the usefulness of our platform, we performed a comprehensive benchmark of existing and new TAD callers exploring different matrix correction methods, parameter settings and sequencing depths. Users can extend our pipeline by adding more tools as they become available.

**Conclusions:** HiC-bench consists an easy-to-use and extensible platform for comprehensive analysis of Hi-C datasets. We expect that it will facilitate current analyses and help scientists formulate and test new hypotheses in the field of three-dimensional genome organization.

## Background

Nuclear organization is of fundamental importance to gene regulation. Recently, proximity ligation assays have greatly enhanced our understanding of chromatin organization and its relationship to gene expression [1]. Here we focus on Hi-C, a powerful genome-wide chromosome conformation capture variant, which detects genome-wide chromatin interactions [2,3]. In Hi-C, chromatin is cross-linked and DNA is fragmented using restriction enzymes, the interacting fragments are ligated forming hybrids that are then sequenced and mapped back to the genome. Hi-C is a very powerful technique that has led to important discoveries regarding the organizational principles of the genome. More specifically, Hi-C has revealed that the mammalian genome is organized in active and repressed areas (A and B compartments) [2] that are further divided in “meta-TADs” [4], TADs [5] and sub-TADs [6]. TADs consist evolutionarily conserved, megabase-scale, non-overlapping areas with increased frequency of intra-domain compared to inter-domain chromatin interactions [5,7]. Despite the fact that Hi-C is very powerful, it is known to be prone to systematic biases [8–10]. Moreover, as the sequencing costs plummet allowing for increased Hi-C resolution, Hi-C poses formidable challenges to computational analysis in terms of data storage, memory usage and processing speed. Thus, various tools have been recently developed to mitigate biases in Hi-C data and make Hi-C analysis faster and more efficient in terms of resource usage. HiC-Box [11], hiclib [9] and HiC-Pro [12] perform various Hi-C analysis tasks, such as alignment and binning of Hi-C sequencing reads into Hi-C contact matrices, noise reduction and detection of specific DNA-DNA interactions. Hi-Corrector [13] has been developed for noise reduction of Hi-C data, allowing parallelization and effective memory management, whereas Hi-Cpipe [14] offers parallelization options and includes steps for alignment, filtering, quality control, detection of specific interactions and visualization of contact matrices. Other tools that allow parallelization are HiFive [15], HOMER [16] and HiC-Pro [12]. Allele-specific Hi-C contact maps can be generated using HiC-Pro and HiCUP [17] (with SNPsplit [18]). TADbit can be used to map raw reads, create interaction matrices, normalize and correct the matrices, call topological domains and build three-dimensional (3D) models based on the Hi-C matrices [19]. HiCdat performs binning, matrix normalization, integration of other data (e.g. ChIP-seq) and visualization [20]. HIPPIE offers similar functionality with HiCdat and allows detection of specific enhancer-promoter interactions [21]. Other tools mainly focus on visualization of Hi-C data (e.g. Sushi [22] and HiCPlotter [23]). Despite the recent boom in the development of computational methods for Hi-C analysis, most of these tools only focus on certain aspects of the analysis, thus failing to encompass the entire Hi-C data analysis workflow. More importantly, these tools or pipelines are not easily extensible, and, for any given Hi-C task, they do not allow the integration of multiple alternative tools (use of alternative TAD calling methods for example) whose performance could then be qualitatively or quantitatively compared. Available tools do not support comprehensive reporting of the parameters used for each task and they do not enable reproducible computational analysis which is an imperative requirement in the era of big data [24], especially given the complexity of Hi-C analyses. The recently released HiFive is an exception as it offers a Galaxy interface [15]. However, use of Galaxy [25] can become problematic for data-heavy analyses, especially when the remote Galaxy server is used.

To facilitate comprehensive processing, reproducibility, parameter exploration and benchmarking of Hi-C data analyses, we introduce HiC-bench, a data flow platform which is extensible and allows the integration of different task-specific tools. Current and future tools related to Hi-C analysis can be easily incorporated into HiC-bench by implementing simple wrapper scripts. HiC-bench covers all current aspects of a standard Hi-C analysis workflow, including read mapping, filtering, quality control, binning, noise correction and identification of specific interactions (Table 1). Moreover, it integrates multiple alternative tools for performing each task (such as matrix correction tools and TAD-calling algorithms), while at the same time allowing simultaneous exploration of different parameter settings that are propagated from one task to all subsequent tasks in the pipeline. HiC-bench also generates a variety of quality assessment plots and offers other visualization options, such as generating genome browser tracks as well as snapshots using HiCPlotter. We have built this platform with reproducibility in mind, as all tools, versions and parameter settings are recorded throughout the analysis. HiC-bench is released as open-source software and the source code is available on GitHub and Zenodo (for details please refer to “Availability of data and material” section). Our team provides installation and usage support.

## Implementation

### The HiC-bench workflow

HiC-bench is a comprehensive computational pipeline for Hi-C sequencing data analysis. It covers all aspects of Hi-C data analysis, ranging from alignment of raw reads to boundary-score calculation, TAD calling, boundary detection, annotation of specific interactions and enrichment analysis. Thus, HiC-bench consists the most complete computational Hi-C analysis pipeline to date (Table 1). Importantly, every step of the pipeline includes summary statistics (when applicable) and direct comparative visualization of the results. This feature is essential for quality control and facilitates troubleshooting. The HiC-bench workflow (Figure 1) starts with the alignment of Hi-C sequencing reads and ends with the annotation and enrichment of specific interactions. More specifically, in the first step, the raw reads (fastq files) are aligned to the reference genome using Bowtie2 [26] (*align*). The aligned reads are further filtered in order to determine those Hi-C read pairs that will be used for downstream analysis (*filter*). A detailed statistics report showing the numbers and percentages of reads assigned to the different categories is automatically generated in the next step (*filter-stats*). The reads that satisfy the filtering criteria are used for the creation of Hi-C contact matrices (*matrix-filtered*). These contact matrices can either be directly visualized in the WashU Epigenome Browser [27] as Hi-C tracks (*tracks*), or further processed using three alternative matrix correction methods: (a) matrix scaling (*matrix-prep*), (b) iterative correction (*matrix-ic*) [9] and (c) HiCNorm (*matrix-hicnorm*) [28]. As quality control, plots of the average number of Hi-C interactions as a function of the distance between the interacting loci are automatically generated in the next step (*matrix-stats*). The Hi-C matrices, before and after matrix correction, are used as inputs in various subsequent pipeline tasks. First, they are directly compared in terms of Pearson or Spearman correlation (*compare-matrices* and *compare-matrices-stats*) in order to estimate the similarity between Hi-C samples. Second, they are used for the calculation of boundary scores (*boundary-scores* and *boundary-scores-pca*), identification of topological domains (*domains*) and comparison of boundaries (*compare-boundaries* and *compare-boundaries-stats*). Third, high-resolution Hi-C matrices are used for detection and annotation of specific chromatin interactions (*interactions* and *annotations*), enrichment analysis in transcription factors, chromatin marks or other segmented data (*annotation-stats*) and visualization of chromatin interactions in certain genomic loci of interest (*hicplotter*). We should note here that HiC-bench is totally extensible and customizable as new tools can be easily integrated into the HiC-bench workflow (see User Manual for more details). In addition to the multiple alternative tools that can be used to perform certain tasks, HiC-bench allows simultaneous exploration of different parameter settings that are propagated from one task to all subsequent tasks in the pipeline (for details please refer to “Main concepts and pipeline architecture” section). For example, after contact matrices are generated and corrected using alternative methods, HiC-bench proceeds with TAD calling using all computed matrices as inputs (Figure 1 and Figure 2A). This unique feature enables the design and execution of complex benchmark studies that may include combinations of multiple tool/parameter choices in each step. HiC-bench focuses on the reproducibility of the analysis by keeping records of the source code, tool versions and parameter settings, and it is the only HiC-analysis pipeline that allows combinatorial parameter exploration facilitating benchmarking of Hi-C analyses.

**Figure 1.**
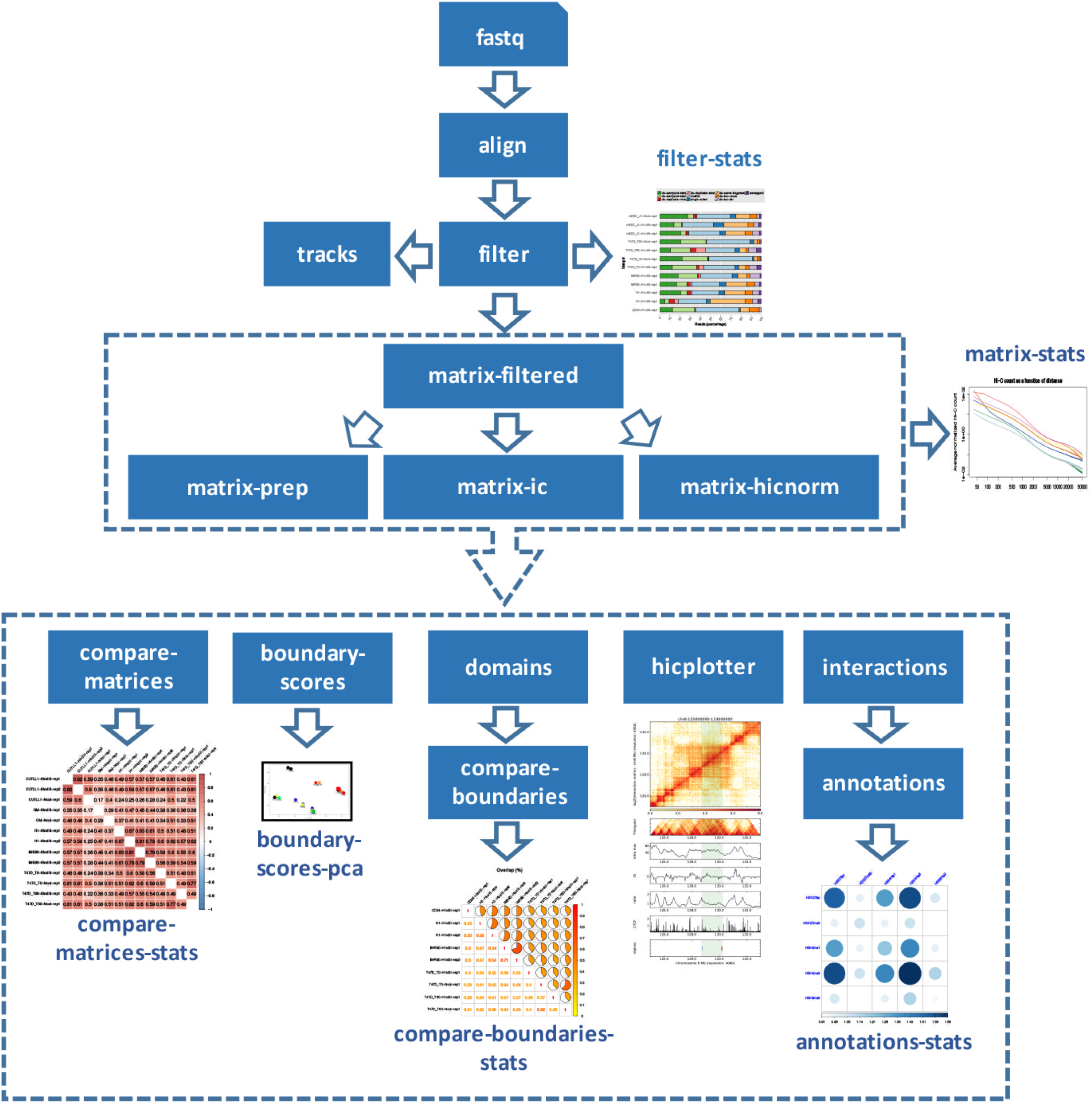
HiC-bench workflow. Raw reads (input fastq files) are aligned and then filtered (*align* and *filter* tasks). Filtered reads are used for the creation of Hi-C track files (*tracks*) that can be directly uploaded to the WashU Epigenome Browser [27]. A report with a statistics summary of filtered Hi-C reads, is also automatically generated (*filter-stats*). Raw Hi-C matrices (*matrix-filtered*) are normalized using (a) scaling, (b) iterative correction [9] or (c) HiCNorm [28]. A report with the plots of the normalized Hi-C counts as function of the distance between the interacting partners (*matrix-stats*) is automatically generated for all methods. The resulting matrices are compared across all samples in terms of Pearson and Spearman correlation (*compare-matrices* and *compare-matrices-stats*). Boundary scores are calculated and the corresponding report with the Principal Component Analysis (PCA) is automatically generated (*boundary-scores* and *boundary-scores-pca*). Domains are identified using various TAD calling algorithms (*domains*) followed by comparison of TAD boundaries (*compare-boundaries* and *compare-boundaries-stats*). A report with the statistics of boundary comparison is also automatically generated. Hi-C visualization of user-defined genomic regions is performed using HiCPlotter (*hicplotter*) [23]. Specific chromatin interactions (*interactions*) are detected and annotated (*annotations*). Finally, enrichment of top interactions in certain chromatin marks, transcription factors etc. provided by the user, is automatically calculated (*annotations-stats*).

**Table 1.** Comparison of HiC-bench with published Hi-C analysis or visualization tools. HiC-bench is a comprehensive and feature-rich Hi-C analysis pipeline that performs various Hi-C tasks by combining our newly-developed tools with existing tools.

| Hi-C tasks | HiC-bench | HiFive | Hi-Cpipe | HiCNorm | hiclib | HiTC | HOMER | Hi-Corrector | HiC-Pro | TADbit | HiCUP | HiC-Box | HiCdat | HIPPIE | Sushi | HiCPlotter |
| --- | --- | --- | --- | --- | --- | --- | --- | --- | --- | --- | --- | --- | --- | --- | --- | --- |
| alignment | x |  | x |  | x |  |  |  | x | x | x | x |  | x |  |  |
| filtering | x | x | x |  | x |  |  |  | x | x | x | x |  | x |  |  |
| genome browser tracks | x |  |  |  |  |  |  |  |  |  |  |  |  |  |  |  |
| quality assessment plots | x | x | x |  |  | x |  |  | x |  | x |  | x | x |  |  |
| contact matrices | x | x | x |  | x |  | x |  | x |  |  | x | x |  |  |  |
| matrix correction | x | x |  | x | x | x | x | x | x | x |  | x | x |  |  |  |
| matrix comparison | x |  |  |  |  |  |  |  |  |  |  |  | x |  |  |  |
| boundary scores | x |  |  |  |  |  |  |  |  |  |  |  |  |  |  |  |
| domains | x |  |  |  |  |  |  |  |  | x |  |  |  |  |  |  |
| boundary comparison | x |  |  |  |  |  |  |  |  |  |  |  |  |  |  |  |
| specific interactions | x |  | x |  | x |  | x |  | x |  | x | x | x | x |  |  |
| annotations | x |  |  |  |  | x |  |  |  |  |  |  | x |  |  |  |
| allele-specific interactions |  |  |  |  |  |  |  |  | x |  | x |  |  |  |  |  |
| visualization | x | x | x |  |  | x | x |  |  |  |  |  | x |  | x | x |
| integration with ChIP-seq data | x |  |  |  |  |  |  |  |  |  |  |  | x | x |  |  |
| parallelization | x | x | x |  |  |  | x | x | x | x |  |  |  |  |  |  |
| integration of alternative tools | x |  |  |  |  |  |  |  |  |  |  |  |  |  |  |  |
| parameter exploration | x |  |  |  |  |  |  |  |  |  |  |  |  |  |  |  |
| reproducibility | x | x |  |  |  |  |  |  |  |  |  |  |  |  |  |  |

### The HiC-bench toolkit

HiC-bench performs various tasks of Hi-C analysis ranging from read alignment to annotation of specific interactions and visualization. We have developed two new tools, *gtools-hic* and *hic-matrix*, to execute the multiple tasks in the HiC-bench pipeline, but we have also integrated existing tools to allow comparative and complementary analyses and facilitate benchmarking. More specifically, the alignment task is performed either with Bowtie2 [26] or with the “align” function of *gtools-hic*, our newest addition to GenomicTools [29]. Likewise, filtering, creation of Hi-C tracks and generation of Hi-C contact matrices are performed using the functions “filter”, “bin/convert” and “matrix” of *gtools-hic* respectively. For advanced users, we have implemented a series of novel features for these common Hi-C analysis tasks. For example, the operation “matrix” of *gtools-hic* allows generation of arbitrary chimeric Hi-C contact matrices, a feature particularly useful for the study of the effect of chromosomal translocations on chromatin interactions. Another example is the generation of distance-restricted matrices (up to some maximum distance off the diagonal) in order to save storage space and reduce memory usage at fine resolutions. For matrix correction we use either published algorithms (iterative correction (IC/ICE) [9], HiCNorm [28]) or our “naïve scaling” method where we divide the Hi-C counts by (a) the total number of (usable) reads, and (b) the “effective length” [8,28] of each genomic bin. We also integrated published TAD callers like DI [5], Armatus [30], TopDom [31], insulation index (Crane) [32] and our own TAD calling method (similar but not identical to contrast index [33,34]) implemented as the “domains” operation in *hic-matrix*. Additionally, the “domains” operation produces genome-wide boundary scores using multiple methods and allowing flexibility in choosing parameters. Boundaries are simply defined as local maxima of the boundary scores. For the detection of specific interactions, we introduce the “loops” function of *hic-matrix*, while GenomicTools is used for annotation of these interactions with gene names, ChIP-seq and other user-defined data. Finally, we implemented a wrapper for HiCPlotter, taking advantage of its advanced visualization features in order to allow the user to quickly generate snapshots of areas of interest in batch. The HiC-bench toolkit is summarized in Table 2. All the tools we developed appear in bold. Further information on the toolkit is provided in the User Manual found online and in the Supplemental Information section.

**Table 2.** The HiC-bench toolkit. The HiC-bench toolkit consists mostly of newly-developed tools (shown in bold) but we have also incorporated existing tools to allow comparisons and benchmarking.

| Hi-C tasks | HiC-bench toolkit |
| --- | --- |
| alignment | bowtie2, **gtools-hic[align]** |
| filtering | **gtools-hic[filter]** |
| genome browser tracks | **gtools-hic[bin/convert]** |
| matrix generation | **gtools-hic[matrix]** |
| matrix correction | IC, HiCNorm, **hic-matrix[preprocess/normalize]** |
| boundary scores | **hic-matrix[domains]** |
| domain calling | DI, Armatus, TopDom, **hic-matrix[domains]** |
| interactions | **hic-matrix[loops]** |
| annotations | **genomic-tools** |
| visualization | HiCPlotter |

### Main concepts and pipeline architecture

We built our platform based on principles outlined in scientific workflow systems such as Kepler [35], Taverna [36] and VisTrails [37]. The main idea behind our platform is the ability to track data provenance [37,38], the origin of the data, computational tasks, tool versions and parameter settings used in order to generate a certain output (or collection of outputs) from a given input (or collection of inputs). Thus, our pipeline ensures reproducibility which is a particularly important feature for such a complex computational task. In addition, HiC-bench enables combinatorial analysis and parameter exploration by implementing the idea of computational “trails”: a unique combination of inputs, tools and parameter values can be imagined as a unique (computational) trail that is followed simultaneously with all the other possible trails in order to generate a collection of output objects (Figure 2A). Our platform consists of three main components: (a) data, (b) code and (c) pipelines. These components are organized in respective directories in our local repository, and synchronized with a remote GitHub repository for public access. The data directory is used to store data that would be used by any analysis, for example genome-related data, such as DNA sequences and indices (e.g. Bowtie2), gene annotations and, in general, any type of data that is required for the analysis. The code directory is used to store scripts, source code and executables. More details about the directory structure can be found in the User Manual. Finally, the “pipelines” directory is used to store the structure of each pipeline. Here, we will focus on our Hi-C pipeline, but we have also implemented a ChIP-seq pipeline, which is very useful for integrating CTCF and histone modification ChIP-seq data with Hi-C data. The structure of the pipeline is presented to the user as a numbered list of directories, each one corresponding to one operation (or task) of the pipeline. As shown in Figure 1, our Hi-C pipeline currently consists of several tasks starting with alignment and reaching completion with the identification and annotation of specific DNA-DNA interactions and annotations with ChIP-seq and other genome-wide data (see also Table 2 and **Supplemental Table 1**). We will examine these tasks in detail in the Results section of this manuscript.

**Figure 2.**
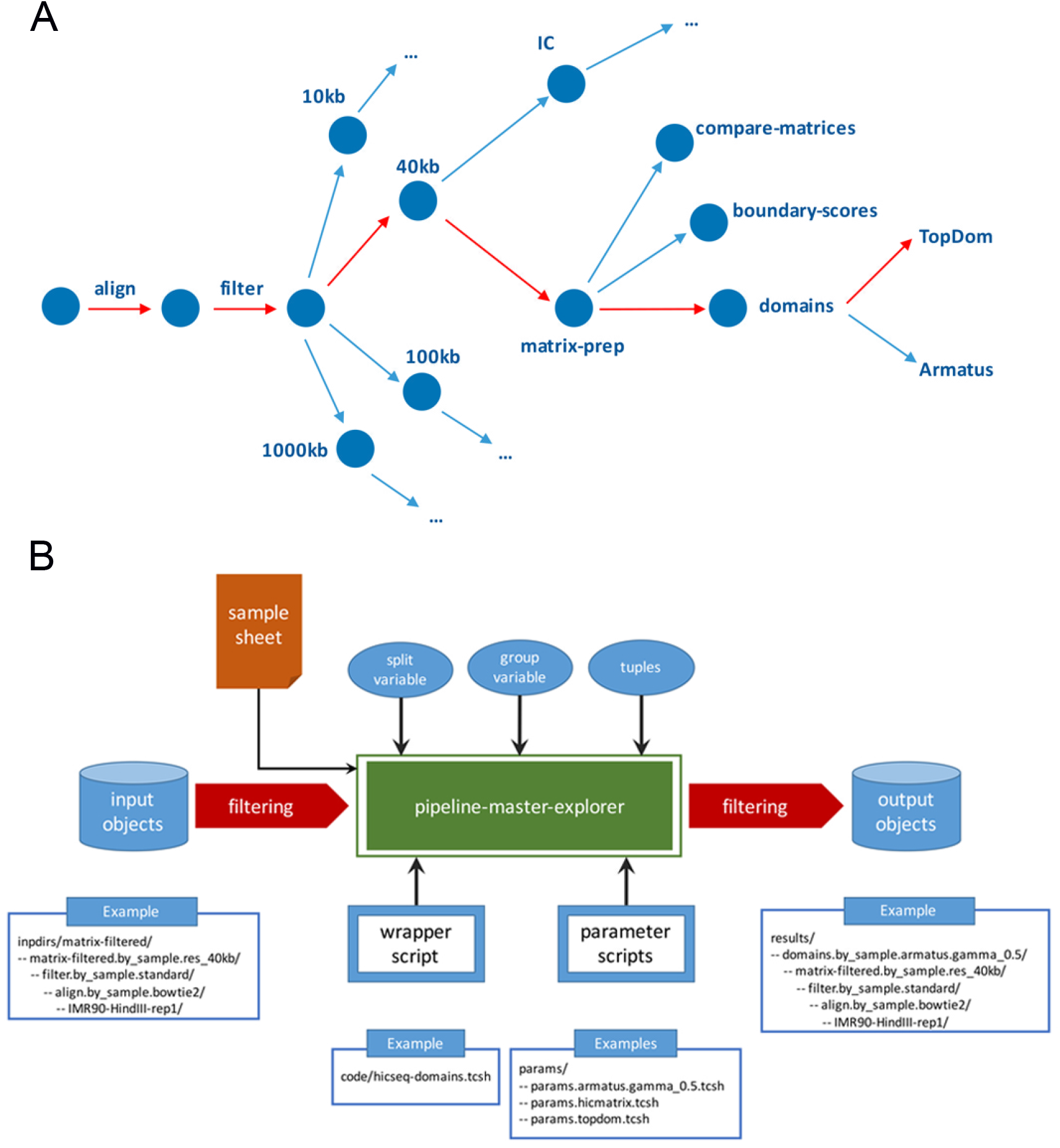
**(A) Computational trails.** Each combination of tools and parameter settings can be imagined as a unique computational “trail” that is executed simultaneously with all the other possible trails to create a collection of output objects. As an example, one of these possible trails is presented in red. The raw reads were aligned, filtered and then binned in 40kb resolution matrices. Our own naïve matrix scaling method was then used for matrix correction and domains were called using TopDom [31]. **(B) HiC-bench pipeline task architecture.** All pipeline tasks are performed by a single R script, “*pipeline-master-explorer.r*”. This script generates output objects based on all combinations of input objects and parameter scripts while taking into account the split variable, group variable and tuple settings. The output objects are stored in the corresponding “*results*” directory. As an example, domain calling for IMR90 is presented. The filtered reads of the IMR90 Hi-C sample (digested with HindIII) are used as input. The pipeline-master-explorer script tests if TAD calling with these settings has been performed and if not it calls the domain calling wrapper script (*code/hicseq-domains.tcsh*) with the corresponding parameters (e.g. *params/params.armatus.gamma_0.5.tcsh*). After the task is complete, the output is stored in the corresponding “results” directory.

### Parameter exploration, input and output objects

In conventional computational pipelines, several computational tasks (operations) are executed on their required inputs. However, in existing genomics pipelines, each task generates a single result object (e.g. TAD calling using one method with fixed parameter settings) which is then used by downstream tasks. To allow full parameter (and method/tool) exploration, we introduce instead a data flow model, where every task may accommodate an arbitrary number of output objects. Downstream tasks will then operate on all computed objects generated by the tasks they depend on. Pipeline tasks are implemented as shown in the diagram of Figure 2B. First, input objects are filtered according to user-specified criteria (e.g. TAD calling is only done for Hi-C contact matrices at 40kb resolution). Then, *pipeline-master-explorer* (implemented as an R script; see User Manual for usage and input arguments) generates the commands that create all desired output objects. In principle, all combinations of input objects with all parameter settings will be created, subject to user-defined filtering criteria. In the interest of extensibility, new pipeline tasks can be conveniently implemented using a single-line *pipeline-master-explorer* command (see **Supplemental Table 2**), provided that wrapper scripts for each task (e.g. TAD calling using TopDom) have been properly set up. In the simplest scenario, any task in our pipeline will generate computational objects for each combination of parameter file and input objects obtained from upstream tasks. For example, suppose the aligned reads from 12 Hi-C datasets are filtered using three different parameter settings, and that we need to create contact matrices at four resolutions (1Mb, 100kb, 40kb and 10kb). Then, the number of output objects (contact matrices in this case) will be 144 (i.e. 12 × 3 × 4). Although many computational scenarios can be realized by this simple one-to-one mapping of input-output objects, more complex scenarios are frequently encountered, as described in the next section.

### Filtering, splitting and grouping input objects into new output objects

Oftentimes, a simple one-to-one mapping of input objects to output objects is not desirable. For this reason, we introduce the concepts of filtering, splitting and grouping of input objects which are used to modify the behavior of pipeline-master-explorer (see Figure 2B). *Filtering* is required when some input objects are not relevant for a given task, e.g. TAD calling is not performed on 1Mb-resolution contact matrices, and specific DNA-DNA interactions are not meaningful for resolutions greater than 10-20kb. *Splitting* is necessary in some cases: for example, we split the input objects by genome assembly (hg19, mm10) when comparing contact matrices or domains across samples, since only matrices or domains from the same genome assembly can be compared directly. In our platform, the user is allowed to split a collection of input objects by any variable contained in the sample sheet (except fastq files), thus allowing user-defined splits of the data, such as by cell type or treatment. Complementary to the splitting concept, *grouping* permits the aggregation of a collection of input objects (sharing the same value of a variable defined in the sample sheet) into a single output object. For example, the user may want to create genome browser tracks or contact matrices of combined technical and/or biological replicates, or group all input objects (samples) together in tasks such as Principal Component Analysis (PCA) or alignment/filtering statistics.

### Combinatorial objects

Even after introducing the concepts described above, more complex scenarios are possible as some tasks require the input of pairs (or triplets etc.) of objects. This feature has also been implemented in our pipeline (tuples in Figure 2B) and is currently used in the *compare-matrices* and *compare-boundaries* tasks. However, it should be utilized wisely (for example in conjunction with filtering, splitting and grouping) because it may lead to a combinatorial “explosion” of output objects.

### Parameter scripts

The design of our platform is motivated by the need to facilitate the use of different parameter settings for each pipeline task. For this reason, we have implemented wrapper scripts for each tool/method used in each task. For example, we have implemented a wrapper script for alignment, filtering, correcting contact matrices using IC or HiCNorm (separate wrappers), TAD calling using Armatus [30], TopDom [31], DI [5] and insulation index (Crane) [32] (separate wrappers). The main motivation is to hide most of the complexity inside the wrapper script and allow the user to modify the parameters using a simple but flexible parameter script. Unlike static parameter files, parameter scripts allow for dynamic calculation of parameters based on certain input variables (e.g. enzyme name, group name etc.). Within this framework, by adding and/or modifying simple parameter scripts, the user can explore the effect of different parameters (a) on the task directly affected by these parameters, and (b) on all dependent downstream tasks. Additionally, these parameter scripts serve as a record of parameters and tool versions that were used to produce the results, facilitating analysis reproducibility as well as documentation in scientific reports and manuscripts.

### Results stored as computational trails

All the concepts described above have been implemented in a single R script named *pipeline-master-explorer*. This script maintains a database of input-output objects for each task, stored in a hidden directory under results (results/.db). It also creates a “run” script which is executed in order to generate all the desired results. All results are stored in the results directory in a tree structure that reveals the computational trail for each object (see examples shown in Figure 2B and **Supplementary Table 2**). Therefore, the user can easily infer how each object was created, including what inputs and what parameters were used.

### Initiating a new reproducible analysis

In the interest of data analysis reproducibility, any new analysis requires creating a copy of the code and pipeline structure into a desired location, effectively creating a branch. This way, any changes in the code repository will not affect the analysis and conversely, the user can customize the code according to the requirements of each project without modifying the code repository. Copying of the code and initiating a new analysis is done simply by invoking the script “*pipeline-new-analysis.tcsh*” as described in the User Manual.

### Pipeline tasks

A pipeline consists of a number of (partially) ordered tasks that can be described by a directed acyclic graph which defines all dependencies. HiC-bench implements a total of 20 tasks as shown in the workflow of Figure 1. In the analysis directory structure, each task is assigned its own subdirectory found inside the pipeline directory starting from the top level. This directory includes a symbolic link to the inputs of the analysis (fastq files, sample sheet, etc.), a link to the code, a directory (*inpdirs*) containing links to all dependencies, a directory containing parameter scripts (see below) and a “*run*” script which can be used to generate all the results of this task. The “*run*” scripts of each task are executed in the specified order by the master “*run*” script located at the top level (see User Manual for details on pipeline directory structure).

### Input data and the sample sheet

Before performing any analysis, a computational pipeline needs input data. All input data for our pipeline tasks are stored in their own “*inputs*” directory accessible at the top level (along with the numbered pipeline tasks) and via symbolic links from within the directories assigned to each task to allow easy access to the corresponding input data. A “*readme*” file explains how to organize the input data inside the inputs directory (see User Manual for details). Briefly, the *fastq* subdirectory is used to store all fastq files, organized into one subdirectory per sample. Then, the sample sheet needs to be generated. This can be done automatically using the “*create-sample-sheet.tcsh*” script, but the user can also manually modify and expand the sample sheet with features beyond what is required. Currently required features are the sample name (to be used in all downstream analyses), fastq files (R1 and R2 in separate columns), genome assembly version (e.g. hg19, mm10) and restriction enzyme name (e.g. HindIII, NcoI). Adding more features, such as different group names (e.g. sample, cell type, treatment), allows the user to perform more sophisticated downstream analyses, such as grouping replicates for generating genome browser tracks, or splitting samples by genome assembly to compare boundaries (see previous section on grouping and splitting).

### Executing the pipeline

The entire pipeline can be executed automatically by the “*pipeline-execute.tcsh*” script, as shown below:

**code/code.main/pipeline-execute <project name> <user e-mail address>**

where <project name> will be substituted by the name of the project and <user e-mail address> by the preferred e-mail address of the person who runs the analysis in order to be notified upon completion. The “*pipeline-execute.tcsh*” script essentially executes the “run” script for each task (following the specified order). At the completion of every task, the log files of all finished jobs are inspected for error messages. If error messages are found, the pipeline aborts with an error message.

### Timestamping

Besides creating the “*run*” script used to generate all results, the “*pipeline-master-explorer.r*” script, also checks whether existing output objects are up-to-date when compared to their dependencies (i.e. input objects and parameter scripts; can be expanded to include code dependencies as well). Currently, the pipelines are set up so that out-of-date objects are not deleted and recomputed automatically, but only presented to the user as a warning. The user can then choose to delete them manually and re-compute. The reason for this is to protect the user against accidentally repeating computationally demanding tasks (e.g. alignments) without given first the chance to review why certain objects may be out-of-date. From a more philosophical point of view, and in the interest of keeping a record of all computations (when possible), the user may never want to modify parameter files or the code for a given project, but instead only add new parameter files. Then, no object will be out-of-date, and only new objects will need to be recomputed every time.

### Alignment and filtering

Paired-end reads were mapped to the reference genome (hg19 or mm10) using Bowtie2 [26]. Reads with low mapping quality (MAPQ<30) were discarded. Local alignments of input read pairs were performed as they consist of chimeric reads between two (non-consecutive) interacting fragments. This approach yielded a high percentage of mappable reads (> 95%) for all datasets (Supplementary Figure 1). Mapped read pairs were subsequently filtered for known artifacts of the Hi-C protocol such as self-ligation, mapping too far from the enzyme’s known cutting sites etc. More specifically, reads mapping in multiple locations on the reference genome (*multihit*), double-sided reads that mapped to the same enzyme fragment (*ds-same-fragment*), reads whose 5’-end mapped too far (*ds-too-far*) from the enzyme cutting site, reads with only one mappable end (*single-sided*) and unmapped reads (*unmapped*), were discarded. Read pairs that corresponded to regions that were very close (less than 25 kilobases, *ds-too-close*) in linear distance on the genome as well as duplicate read pairs (*ds-duplicate-intra* and *ds-duplicate-inter*) were also discarded. In Supplementary Figure 1, we show detailed paired-end read statistics for the Hi-C datasets used in this study. We include the read numbers (Supplementary Figure 1A) and their corresponding percentages (Supplementary Figure 1B). Eventually, approximately 10-50% of paired-reads passed all filtering criteria and were used for downstream analysis (Supplementary Figure 1B). The statistics report is automatically generated for all input samples. The tools and parameter settings used for the alignment and filtering tasks are fully customizable and can be defined in the corresponding parameter files.

### Contact matrix generation, normalization and correction

The read-pairs that passed the filtering task were used to create Hi-C contact matrices for all samples. The elements of each contact matrix correspond to pairs of genomic “bins”. The value in each matrix element is the number of read pairs aligning to the corresponding genomic regions. In this study, we used various resolutions, ranging from fine (10kb) to coarse (1Mb). The resulting matrices either remained unprocessed (filtered), or they were processed using different correction methods including HiCNorm [28], iterative correction (IC or ICE) [9] as well as “naïve scaling”. In Supplementary Figure 2, we present the average Hi-C count as a function of the distance between the interacting fragments, separately for each Hi-C matrix for not corrected (filtered) and IC-corrected matrices.

### Comparison of contact matrices

Our pipeline allows direct comparison and visualization of the generated Hi-C contact matrices. More specifically, using our *hic-matrix* tool, all pairwise Pearson and Spearman correlations were automatically calculated for each (a) input sample, (b) resolution, and (c) matrix correction method. The corresponding correlograms were automatically generated using the corrgram R package [39]. A representative example is shown in Supplementary Figure 3. The correlograms summarizing the pairwise Pearson correlations for all samples used in this study are presented before and after matrix correction using the iterative correction algorithm. These plots are very useful because the user can quickly assess the similarity between technical and biological replicates as well as differences between various cell types. As shown before (Supplementary Figure 3 in [5]), iterative correction improves the correlation between enzymes at the expense of a decreased correlation between samples prepared using the same enzyme.

### Boundary scores

Topological domains (TADs) are defined as genomic neighborhoods of highly interacting chromatin, with relatively more infrequent inter-domain interactions [5,40,41]. Topological domains are demarcated by boundaries, i.e. genomic regions bound by insulators thus hampering DNA contacts across adjacent domains. For each genomic position, in a given resolution (typically 40kb or less), we define a “boundary score” to quantify the insulation strength of this position. The higher the boundary score, the higher the insulation strength and the probability that this region actually acts as a boundary between adjacent domains. The idea of boundary scores is further illustrated in Supplementary Figure 4, where two adjacent TADs are shown. The upstream TAD on the left (*L*) is separated from the downstream TAD on the right (*R*) by a boundary (black circle). We define two parameters, the distance from the diagonal of the Hi-C contact matrix to be excluded from the boundary score calculation (*δ*) (not shown) and the maximum distance from the diagonal to be considered (*d*). The corresponding parameter values can be selected by the user. For this analysis, we used *δ*=0 and *d*=2Mb as suggested before [5]. In addition to the published directionality index [5], we defined and computed the “*inter*”, “*intra-max*” and “*ratio*” scores, defined as follows:

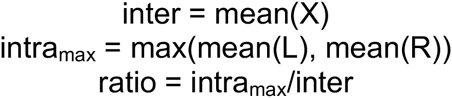

Principal component analysis (PCA) of boundary scores across samples in this study,459460before and after matrix correction, shows that biological replicates tend to cluster461together, either in the case of filtered or corrected (IC) matrices (Supplementary Figure 5).

### Topological domains

Topologically-associated domains (TADs) are increasingly recognized as an important feature of genome organization [5]. Despite the importance of TADs in genome organization, very few Hi-C pipelines have integrated TAD calling (e.g. TADbit [19]). In HiC-bench, we have integrated TAD calling as a pipeline task and we demonstrate this integration using different TAD callers: (a) Armatus [30], (b) TopDom [31], (c) DI [5], (d) insulation index (Crane) [32] and (e) our own hic-matrix (domains). Our pipeline makes it straightforward to plug in additional TAD callers, by installing these tools and setting up the corresponding wrapper scripts. HiC-bench also facilitates the direct comparison of TADs across samples by automatically calculating the number of TAD boundaries and all the pairwise overlaps of TAD boundaries across all inputs, generating the corresponding graphs (as in the case of matrix correlations described in a previous section). We define boundary overlap as the ratio of the intersection of boundaries between two replicates (A and B) over the union of boundaries in these two replicates, as shown in the equation below:

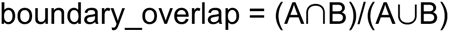

For the boundary overlap calculation, we extended each boundary by 40kb on both sides (+/−40kb flanking region, i.e. the size of one bin). The fact that HiC-bench allows simultaneous exploration of all parameter settings for all installed TAD-calling algorithms, greatly facilitates parameter exploration, optimization as well as assessment of algorithm effectiveness. Pairwise comparison of boundaries (boundary overlaps) across samples is shown in Figure 3 and Supplementary Figure 6.

**Figure 3.**
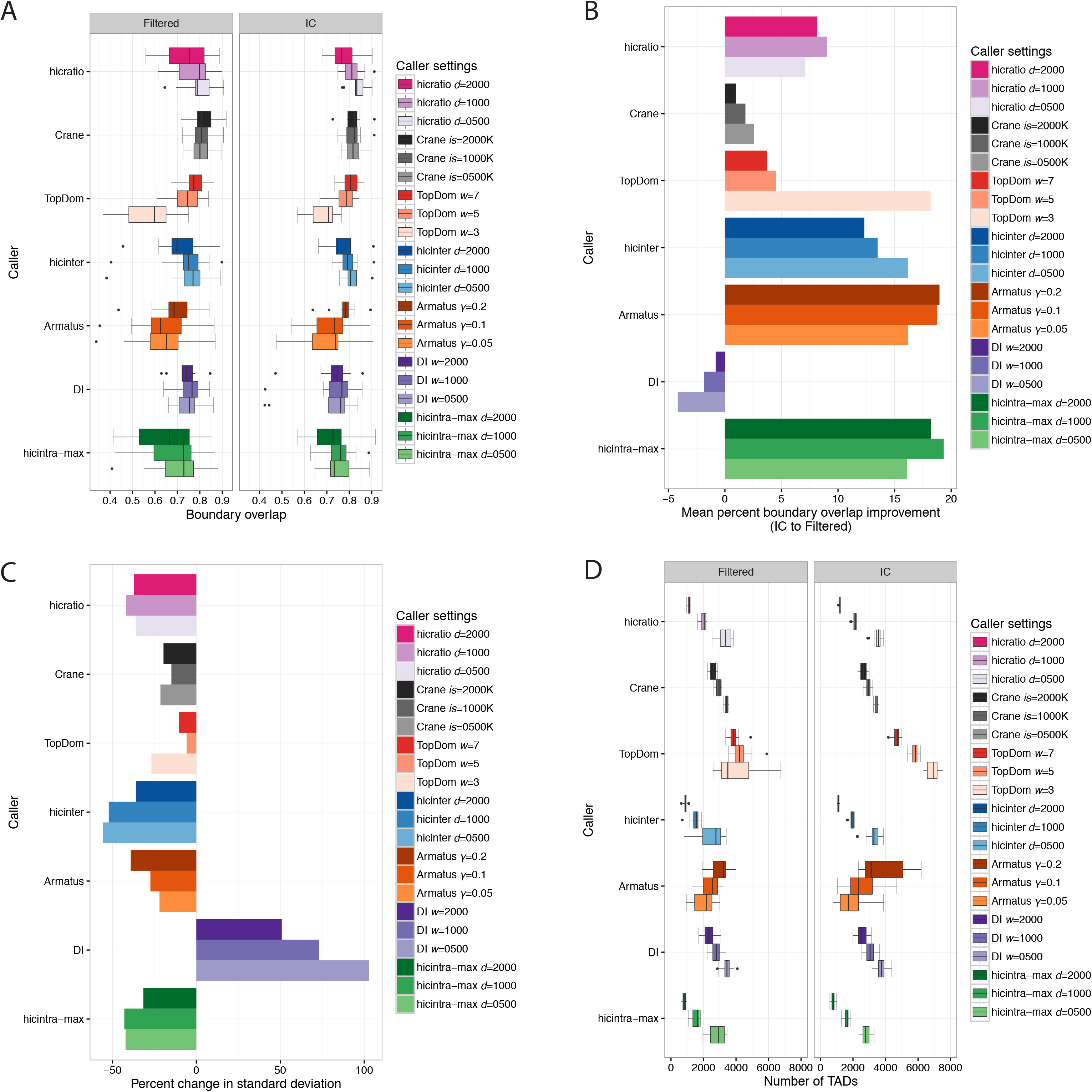
Comparison of topological domain calling methods subject to Hi-C contact matrix preprocessing by simple filtering or iterative correction (IC). The methods were assessed in terms of boundary overlap between replicates (**A**), change (%) in mean boundary overlap after matrix correction (**B**), change (%) in standard deviation of mean overlap across replicates after matrix correction (**C**) and number of identified topological domains per cell type (**D**). The different colors correspond to the different callers. Gradients of the same color are used for the different values of the same parameter, ranging from low (light color) to high (dark color) values. The TAD callers along with the corresponding parameter settings are presented in the legend. For this analysis all available read pairs were used.

### Visualization

In our pipeline, we also take advantage of the great visualization capabilities offered by the recently released HiCPlotter [23], in order to allow the user to visualize Hi-C contact matrices along with TADs (in triangle format) for multiple genomic regions of interest. The user can also add binding profiles in BedGraph format for factors (e.g. CTCF), boundary scores, histone marks of interest (e.g. H3K4me3, H3K27ac) etc. An example is shown in **Supplementary Figure 7**, where an area of the contact matrix of human embryonic stem cells (H1) (HindIII) is presented along with the corresponding TADs (triangles), various boundary scores, the CTCF binding profile and annotations of selected genomic elements, before and after matrix correction (IC). The integration of HiCPlotter in our pipeline, allows the user to easily create publication-quality figures for multiple areas of interest simultaneously.

### Specific interactions, annotations and enrichments

The plummeting costs of next-generation sequencing have resulted in a dramatic increase in the resolution achieved in Hi-C experiments. While the original Hi-C study reported interaction matrices of 1Mb resolution [2], recently 1kb resolution was reported [42]. Thus, the characterization and annotation of specific genomic interactions from Hi-C data is an important feature of a modern Hi-C analysis pipeline. HiC-bench generates a table of the interacting loci based on parameters defined by the user. These parameters include the resolution, the lowest number of read pairs required per interacting area as well as the minimum distance between the interacting partners. The resulting table contains the coordinates of the interacting loci, the raw count of interactions between them, the number of interactions after “scaling” and the number of interactions between the partners after distance normalization (observed Hi-C counts normalized by expected counts as a function of distance). This table is further annotated with the gene names or the factors (e.g. CTCF) and histone modification marks (e.g. H3K4me1, H3K27ac, H3K4me3) that overlap with the interacting loci. The user can provide bed files with the features of interest to be used for annotation. As an example, the enrichment of chromatin marks in the top 50000 chromatin interactions in the H1 and IMR90 samples is presented in Supplementary Figure 8.

### Software requirements

The main software requirements are: Bowtie2 aligner [26], Python (2.7 or later) (along with Numpy, Scipy and Matplotlib libraries), R (3.0.2) [43] and various R packages (lattice, RColorBrewer, corrplot, reshape, gplots, preprocessCore, zoo, reshape2, plotrix, pastecs, boot, optparse, ggplot2, igraph, Matrix, MASS, flsa, VennDiagram, futile.logger and plyr). More details on the versions of the packages can be found in the User Manual (sessionInfo()). In addition, installation of mirnylib Python library [44] is required for matrix balancing based on IC (ICE). The pipeline has been tested on a high-performance computing cluster based on Sun Grid Engine (SGE). The operating system used was Redhat Linux GNU (64 bit).

## Results

We used HiC-bench to analyze several published Hi-C datasets and the results of our analysis are presented below. Additionally, we performed a comprehensive benchmark of existing and new TAD callers exploring different matrix correction methods, parameter settings and sequencing depths. Our results can be reproduced by re-running the corresponding pipeline snapshot available upon request as a single compressed archive file (too big to include as a Supplemental file).

### Comprehensive reanalysis of available Hi-C datasets using HiC-bench

Our platform is designed to facilitate and streamline the analysis of a large number of available Hi-C datasets in batch. Thus, we collected and comprehensively analyzed multiple Hi-C samples from three large studies [5,42,45]. From the first study we analyzed IMR90 (HindIII) samples, from the second we analyzed Hi-C samples from lymphoblastoid cells (GM12878), human lung fibroblasts (IMR90 (MboI)), erythroleukemia cells (K562), chronic myelogenous leukemia (CML) cells (KBM-7) and keratinocytes (NHEK), and from the third one, we analyzed samples from human embryonic stem cells (H1) and all the embryonic stem-cell derived lineages mentioned, including mesendoderm, mesenchymal stem cells, neural progenitor cells and trophectoderm cells. All datasets yielded at least 40 million usable intra-chromosomal read pairs in at least two biological replicates. We performed extensive quality control on all datasets, calculating the read counts and percentages per classification category (Supplementary Figure 1), the attenuation of Hi-C signal over genomic distance (Supplementary Figure 2), the correlation of Hi-C matrices before and after matrix correction (Supplementary Figure 3), the similarity of boundary scores (Supplementary Figure 5) and all pairwise boundary overlaps across samples (Supplementary Figure 6). In addition, we performed a comprehensive benchmarking of our own and published TAD callers, across all reanalyzed Hi-C datasets. The results of our benchmark are presented in the following sections.

### Iterative correction of Hi-C contact matrices improves reproducibility of TAD boundaries

Iterative correction has been shown to correct for known biases in Hi-C [9]. Thus, we hypothesized that IC may increase the reproducibility of TAD calling. We performed a comprehensive analysis calculating the TAD boundary overlaps for biological replicates of the same sample (as described in Methods section), using different TAD callers and different main parameter values for each TAD caller (Figure 3A). After comparing TAD boundary overlaps between filtered (uncorrected) and IC-corrected matrices, we observed an improvement in the boundary overlap when corrected matrices were used, irrespective of TAD caller and parameter settings (Figure 3B). The only exception was DI. Careful examination of the overlaps per sample revealed that IC introduced outliers only in the case of DI (in general, the opposite was true for the other callers). We hypothesize that IC may occasionally negatively affect the computation of the directionality index, especially because its calculation depends on a smaller number of bins (1-dimensional line) compared to the rest of the methods (2-dimensional triangles). In addition to the increase in the mean value of boundary overlap upon correction, we observed that the standard deviation of boundary overlaps among replicates decreased (again, with the exception of DI) (Figure 3C). While this seems to be the trend for almost all TAD caller/parameter value combinations, the effect of correction in variance is more profound in certain cases (e.g. hicintra-max) than others. It is also worth mentioning that increased size of the insulation window (in the case of Crane), the resolution parameter *γ* (Armatus) or the distance *d* (hicinter, hicintra-max, hicratio) may result in certain cases in increased boundary overlap (e.g. Armatus), but this is not generalizable (e.g. hicintra-max). Interestingly, increased TAD boundary overlap does not necessarily mean increased consistency in the number of predicted TADs across sample types, as would be expected since TADs are largely invariant across cell types [5]. For example, the TAD calling algorithm which is based on insulation score (Crane), predicted similar TAD overlaps and similar TAD numbers for different insulation windows (ranging from 0.5Mb to 2Mb), whereas Armatus performed well in terms of TAD boundary reproducibility (Figure 3A) but the corresponding predicted TAD numbers varied considerably (Figure 3D). This may be partly due to the nature of the Armatus algorithm, as it has been built to reveal multiple levels of chromatin organization (TADs, sub-TADs etc.). We conclude that while iterative correction improves the reproducibility of TAD boundary detection across replicates, the number of predicted TADs should be also taken into account when selecting TAD calling method for downstream analysis. The method of choice should be the one that is robust in terms of both reproducibility and number of predicted TADs.

### Increased sequencing depth improves the reproducibility of TAD boundaries

After selecting the parameter setting that optimized TAD boundary overlap between biological replicates of the same sample per TAD caller, we also investigated the effect of sequencing depth on the reproducibility of TAD boundary detection. Since some of the input samples were limited to only 40 million usable intra-chromosomal Hi-C read pairs, we resampled 10 million, 20 million and 40 million read pairs from each sample and evaluated the effect of increasing sequencing depth on TAD boundary reproducibility. The results are depicted in Figure 4A. We noticed that increased sequencing depth resulted in increased TAD boundary overlap, regardless of the TAD calling algorithm used (Figure 4A,C). As far as the TAD numbers are concerned, increased sequencing depth decreased TAD number variability for certain callers (e.g. hicratio) but not in all cases (e.g. Armatus) (Figure 4B). In many cases, increased sequencing depth, decreased the variance of TAD boundary overlap among replicates (Figure 4C). In summary, based on this benchmark, we recommend that Hi-C samples should be sufficiently sequenced as sequencing depth seems to affect TAD calling reproducibility.

**Figure 4.**
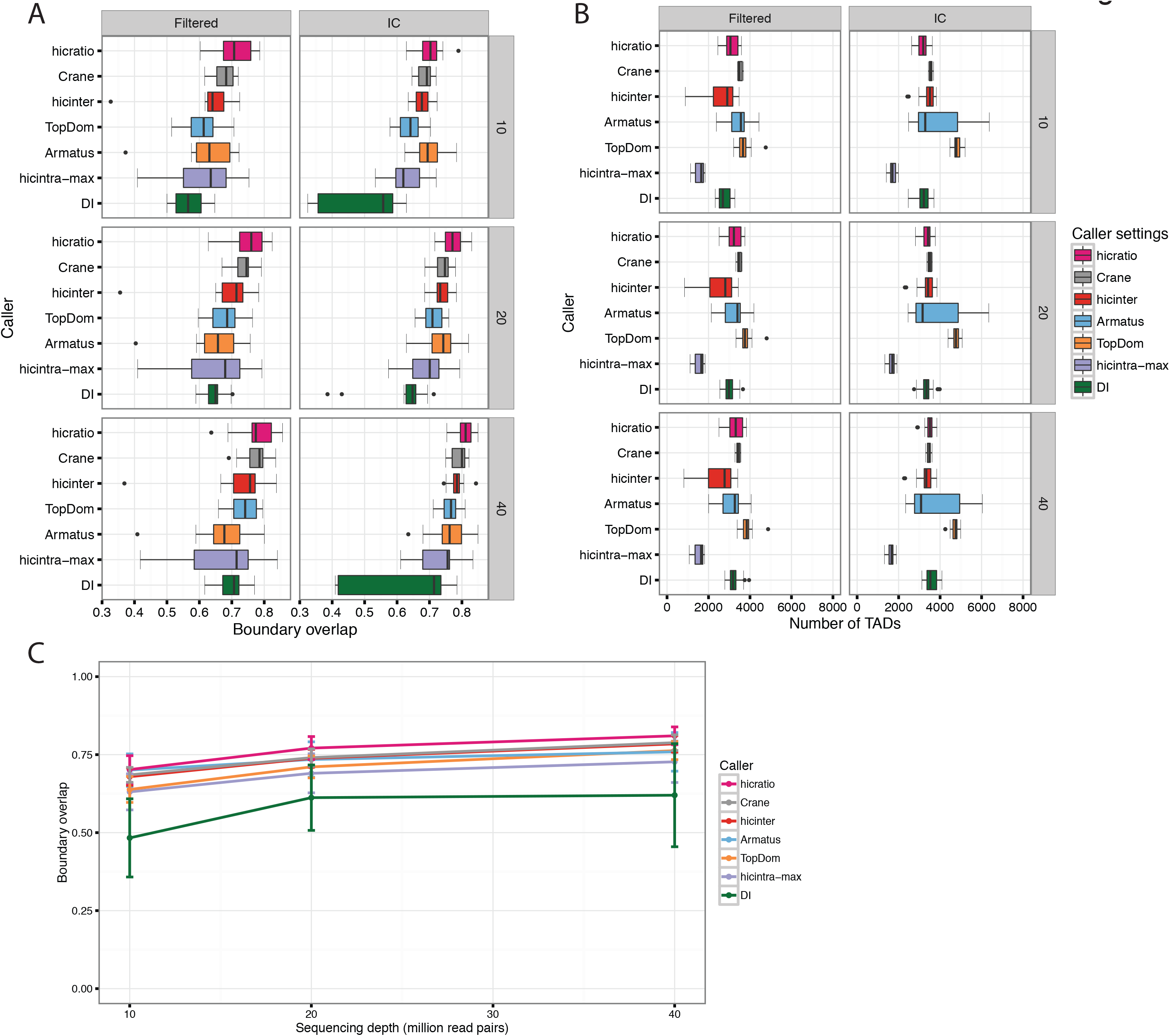
Comparison of topological domain calling methods for different preprocessing method and sequencing depth. TAD calling methods were assessed in terms of boundary overlap between replicates (**A**), number of identified topological domains (**B**) and boundary overlap across replicates upon increasing sequencing depth (**C**) for different matrix preprocessing (filtered and IC corrected) and different sequencing depths (10 million, 20 million and 40 million reads). For TAD calling, only the optimal caller/parameter value pairs are shown (defined as the ones achieving the maximum boundary overlap for IC and 40 million reads). The boxplot and line colors correspond to the different TAD callers.

## Conclusions

Recently, several computational tools and pipelines have been developed for Hi-C analysis. Some focus on matrix correction, others on detection of specific chromatin interactions and their differences across conditions and others on visualization of these interactions. However, very few of these tools offer a complete Hi-C analysis (e.g. HiC-Pro), addressing tasks which range from alignment to interaction annotation. HiC-bench is a comprehensive Hi-C analysis pipeline with the ability to process many samples in parallel, record and visualize the results in each task, thus facilitating troubleshooting and further analyses. It integrates both existing tools but also new tools that we developed to perform certain Hi-C analysis tasks. In addition, HiC-bench focuses on parameter exploration, reproducibility and extensibility. All parameter settings used in each pipeline task are automatically recorded, while future tools can be easily added using the supplied wrapper template. More importantly, HiC-bench is the only Hi-C pipeline so far that allows extensive parameter exploration, thus facilitating direct comparison of the results obtained by different tools, methods and parameters. This unique feature helps users test the robustness of the analysis, optimize the parameter settings and eventually obtain reliable and biologically meaningful results. To demonstrate the usefulness of HiC-bench, we performed a comprehensive benchmark of popular and newly-introduced TAD callers, varying the matrix preprocessing (filtered or corrected matrices with IC method), the sequencing depth, and the value of the main parameter of each TAD caller, which is usually the window used for the calculation of directionality index or insulation score. We found that the matrix correction has a positive effect on the boundary overlap between replicates and that increased sequencing depth leads to higher boundary overlap.

In conclusion, HiC-bench is an easy-to-use framework for systematic, comprehensive, integrative and reproducible analysis of Hi-C datasets. We expect that use of our platform will facilitate current analyses and enable scientists to further develop and test interesting hypotheses in the field of chromatin organization and epigenetics.

## List of abbreviations

DI: directionality index
IC or ICE: iterative correction
PCA: Principal Component Analysis
TAD: Topological Domain or Topologically Associating Domain

## Ethics approval and consent to participate

Not applicable.

## Consent for publication

Not applicable.

## Availability of data and material

Published Hi-C data were downloaded from Gene Expression Omnibus, using the corresponding accession numbers: GSE35156 [5], GSE63525 [42] and GSE52457 [45].

HiC-bench source code is freely available on GitHub and Zenodo.

Project name: HiC-bench

Project home page: <u>https://github.com/NYU-BFX/hic-bench/wiki</u>

Archived version: <u>https://zenodo.org/badge/latestdoi/20915/NYU-BFX/hic-bench</u>

Operating system: Redhat Linux GNU (64 bit)

Programming language: R, C++, Python, Unix shell scripts

Other requirements: mentioned in “Software requirements” section and the manual.

License: MIT

Any restrictions to use by non-academics: None

## Author contributions

CL performed computational analyses, generated figures and implemented certain wrapper scripts. SK wrote the user manual. PN and IA offered biological insights and helped with the interpretation of Hi-C data. AT designed and implemented the pipeline. CL and AT wrote the manuscript.

## Acknowledgements

Aristotelis Tsirigos was supported by a Research Scholar Grant, RSG-15-189-01 - RMC from the American Cancer Society and a Leukemia & Lymphoma Society New Idea Award, 8007-17. We would like to thank Dennis Shasha and Juliana Freire for inspiring discussions on data flows. We would also like to thank Kadir Caner Akdemir for useful discussions on the usage of HiCPlotter. We also like to thank the NYU Applied Bioinformatics Laboratories for providing bioinformatics support and helping with the analysis and interpretation of the data. This work has used computing resources at the High Performance Computing Facility (HPCF) at the NYU Langone Medical Center.

---

**Supplementary Figure 1.**
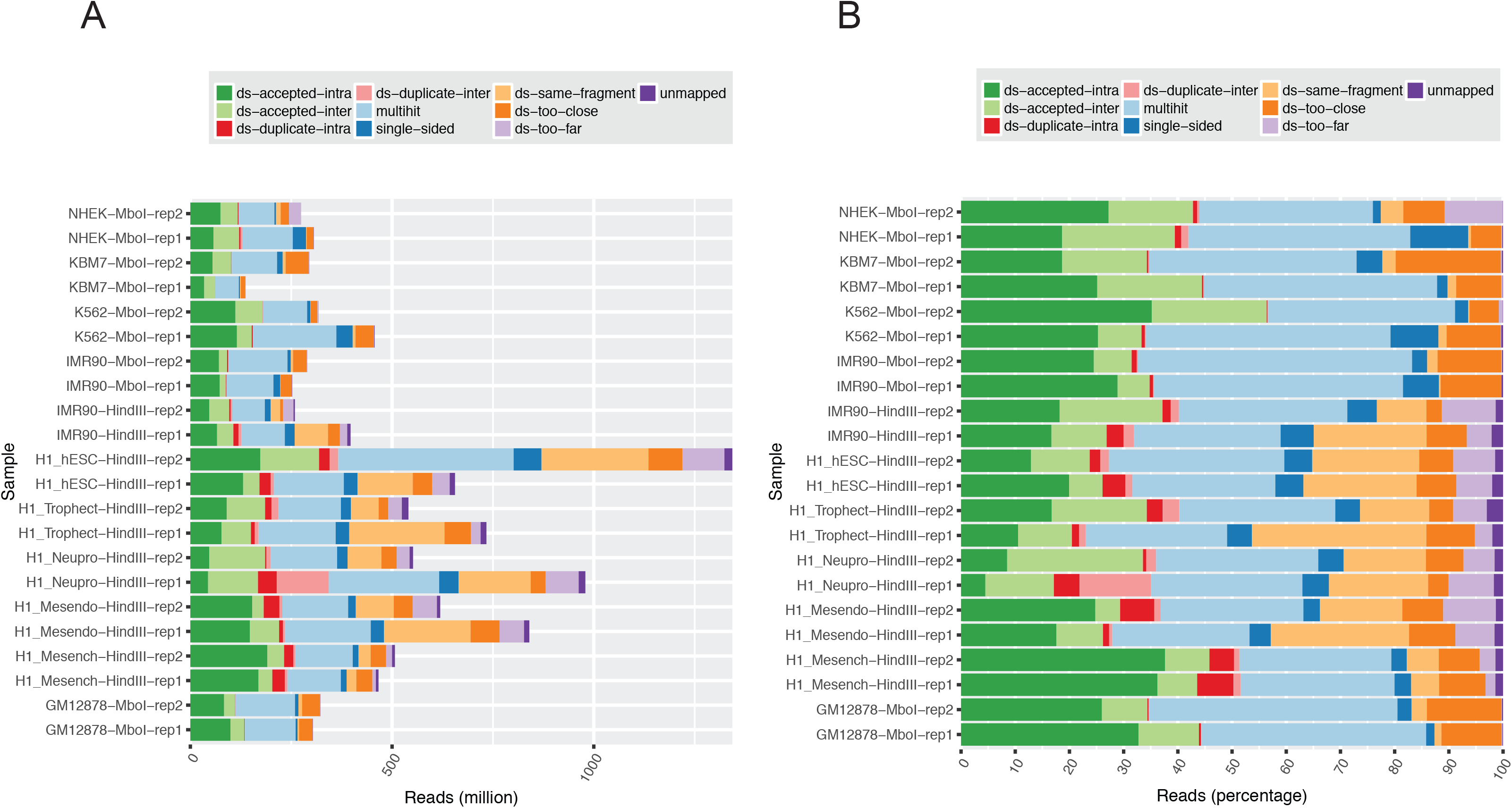
Hi-C reads filtering statistics. Number **(A)** and percentage (**B**) of the various read categories identified during filtering for all datasets used in the study. Mappable reads were over 95% in all samples. Duplicate (*ds-duplicate-intra* and *ds-duplicate-inter*; red and pink), non-uniquely mappable (*multihit*; light blue) and single-end mappable (*single-sided*; dark blue) reads were discarded. Self-ligation products (*ds-same-fragment*) and reads mapping too far (*ds-too-far*; light purple) from restriction sites or too close to one another (*ds-too-close*; orange) were also discarded. Only double-sided uniquely mappable *cis* (*ds-accepted-intra*; dark green) and *trans* (*ds-accepted-inter*; light green) read pairs were used for downstream analysis. The *x* axis represents either the raw read number (A) or the percentage of reads (B) falling within each of the categories described in the legend. The *y* axis represents the samples.

**Supplementary Figure 2.**
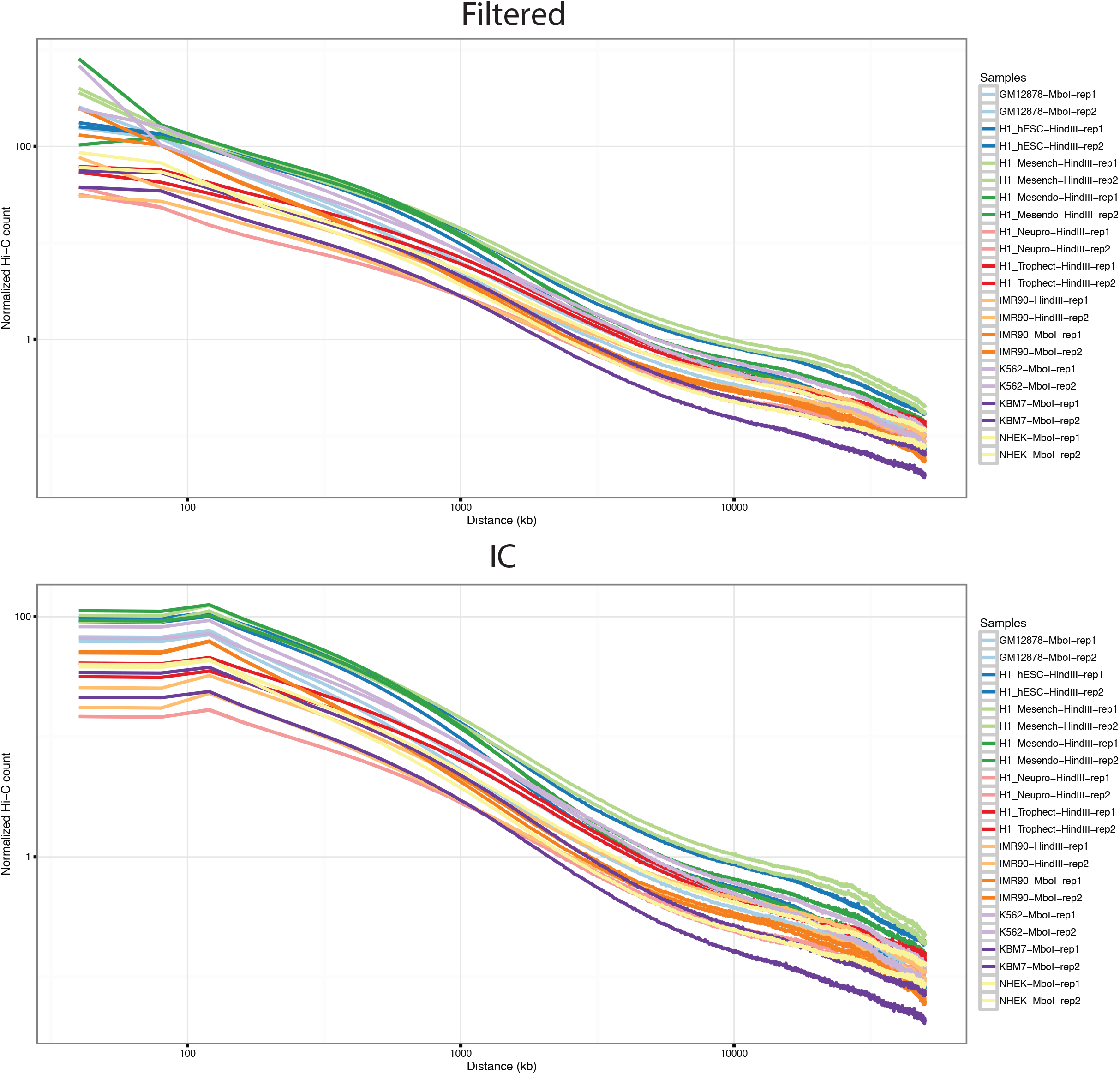
Matrix statistics. Normalized Hi-C counts are presented as a function of the distance between the interacting partners for all samples and correction methods. The Hi-C samples analyzed were GM12878 (light blue), hESCs (H1) (blue), mesenchymal cells (light green), mesendoderm (dark green), neural progenitors (pink), trophectoderm (red), IMR90 (light and dark orange), K562 (light purple), KBM7 (dark purple) and NHEK (yellow). The matrices were either unprocessed (filtered) (top) or corrected using IC (bottom). The *y* axis represents the normalized count of Hi-C interactions and the *x* axis the distance between the interacting partners in kilobases.

**Supplementary Figure 3.**
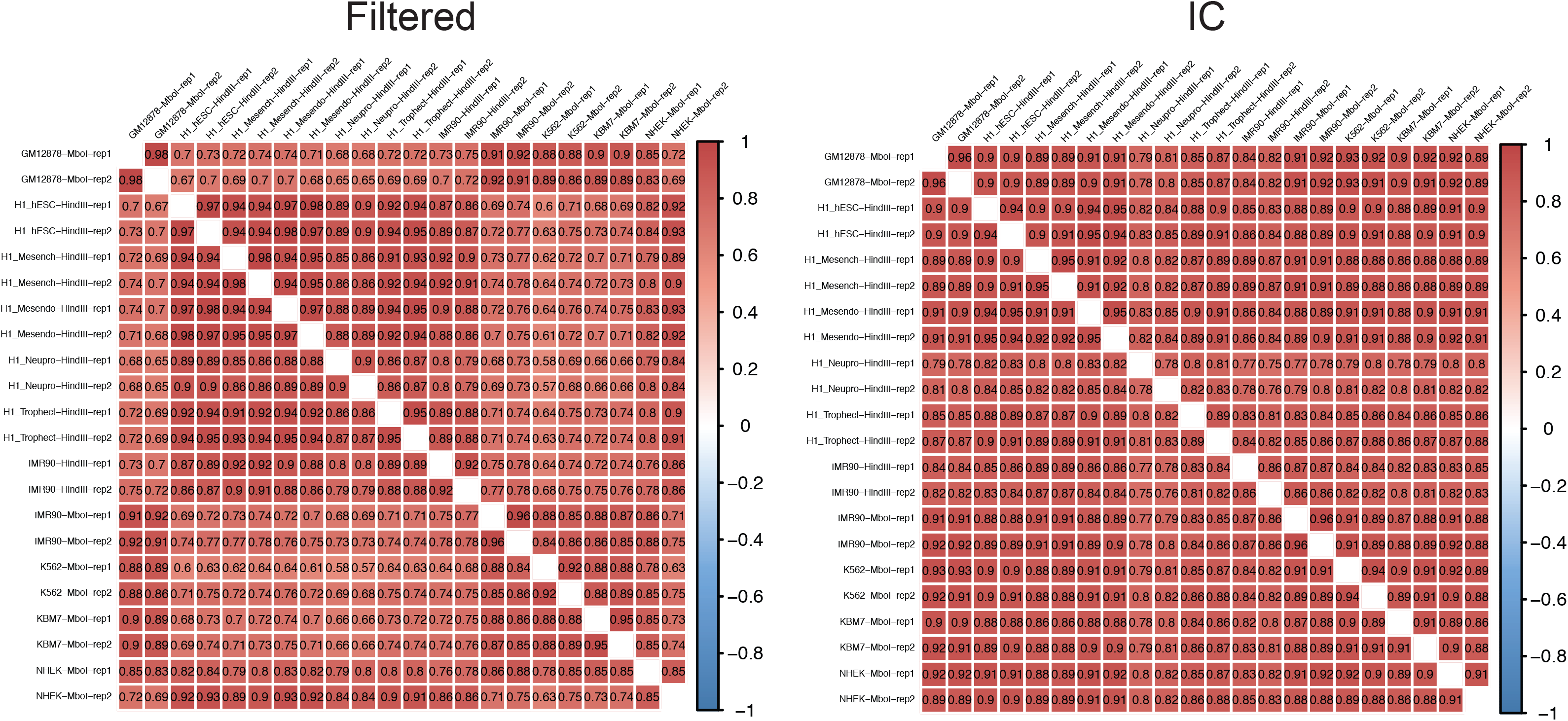
Pairwise Pearson correlation of Hi-C matrices. Correlograms summarizing all pairwise Pearson correlations for all Hi-C samples used in this study: raw (filtered) matrices (left panel) and matrices after iterative correction (right panel). Dark red indicates strong positive correlation and dark blue strong negative. The resolution of the matrices is 40kb.

**Supplementary Figure 4.**
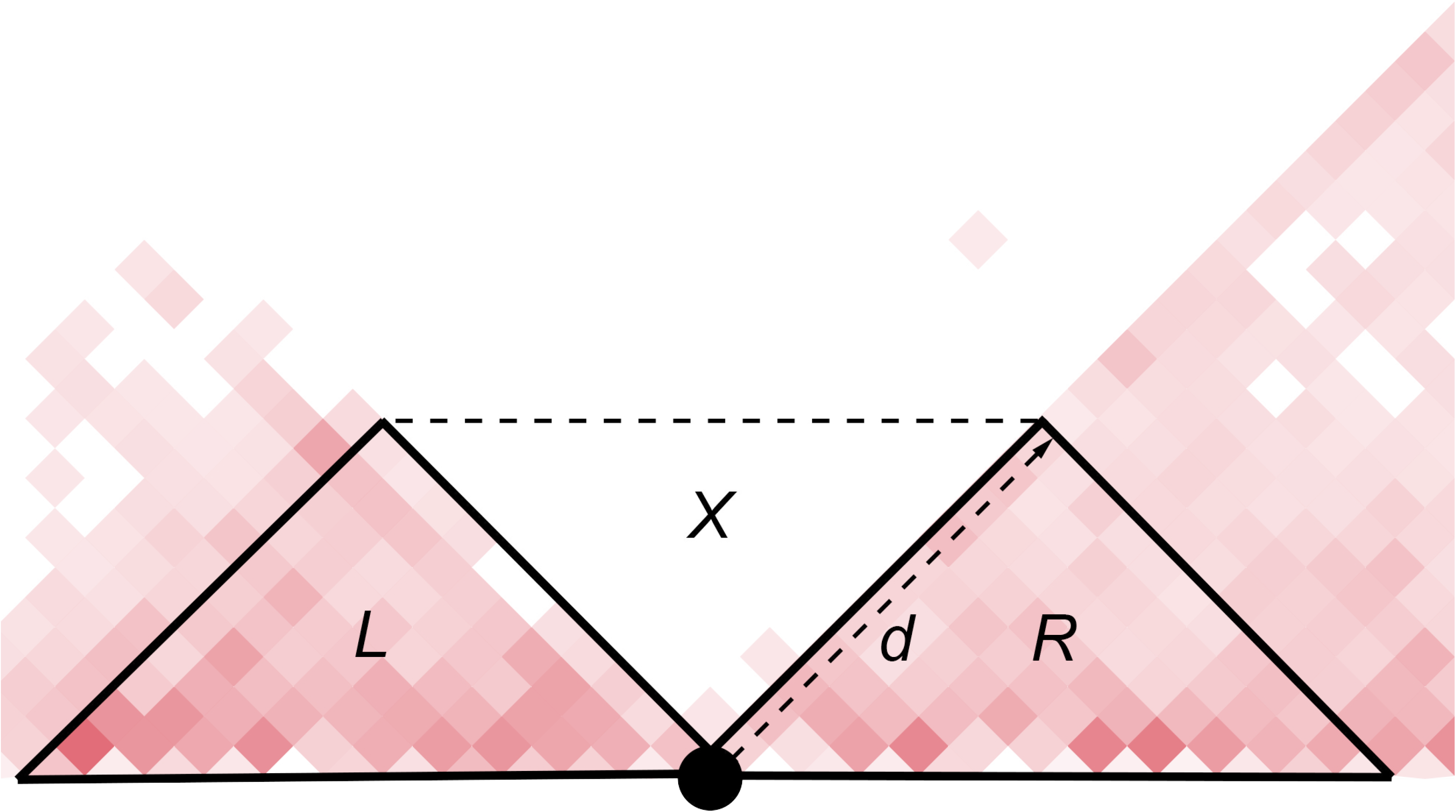
Boundary score calculation. Two adjacent topological domains (red triangles) are depicted. The left domain (*L*) is separated from the right domain (*R*) by a boundary (black circle). The areas of more-frequent intra-domain interactions are in red. The area of less-frequent cross-domain (or inter-domain) interactions is *X*. We also introduce parameter *d* which is the maximum distance from the diagonal to be considered for the calculation of boundary scores (default value: *d*=2Mb).

**Supplementary Figure 5.**
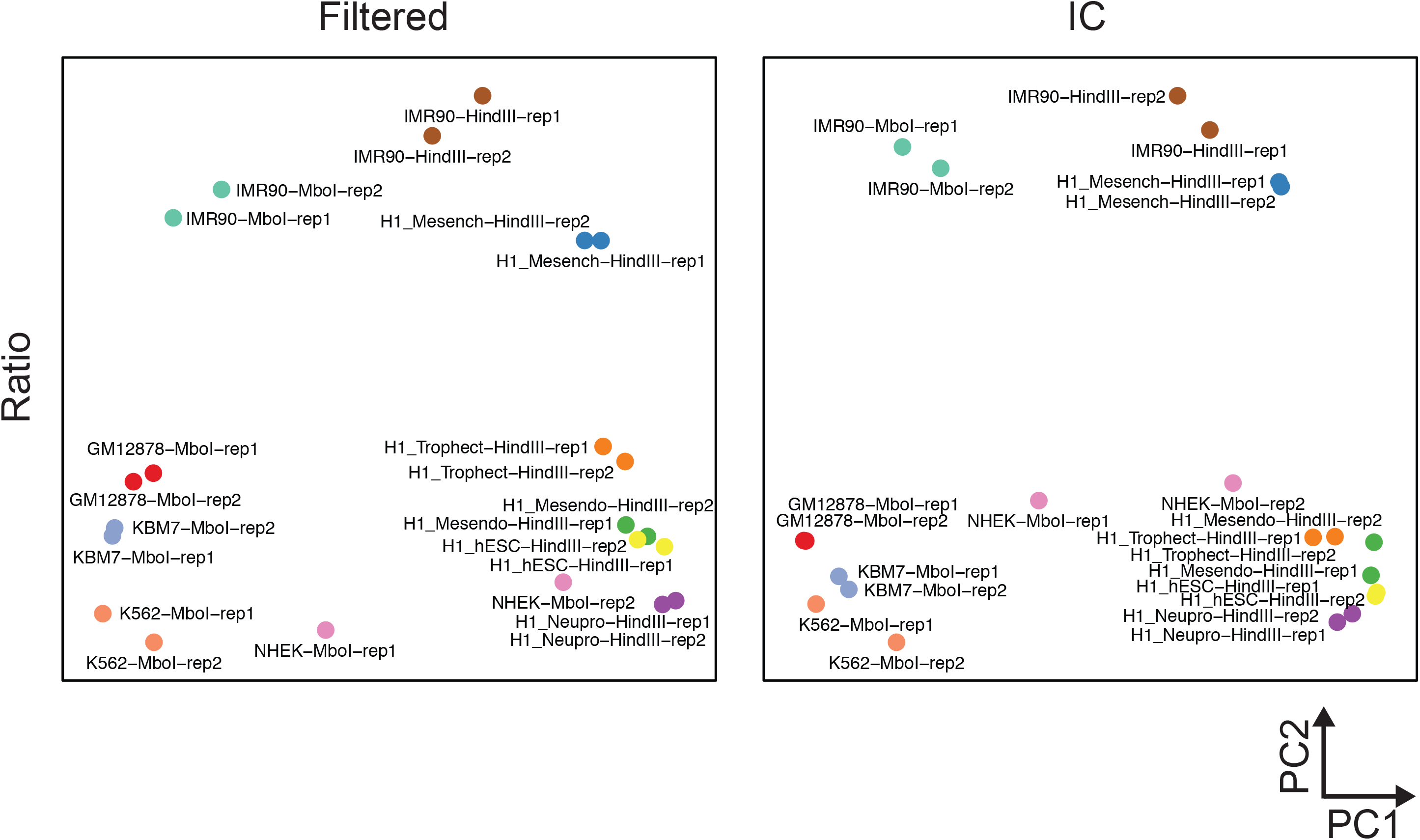
Principal component analysis of boundary scores. Boundary scores were calculated using *ratio* score, for all samples either before (filtered) (left panel) or after iterative correction (IC) (right panel).

**Supplementary Figure 6.**
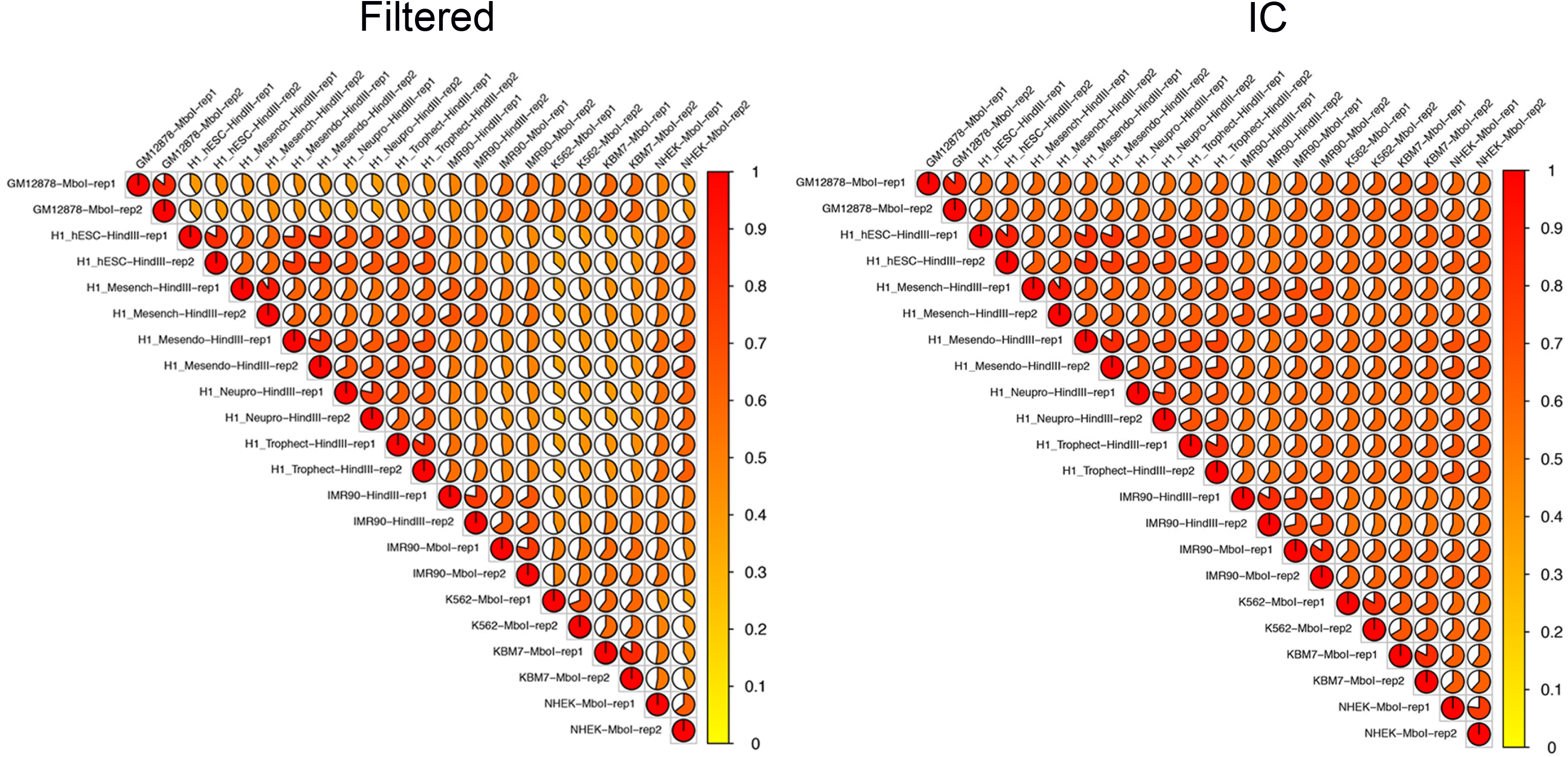
Pairwise overlaps of TAD boundaries. The pairwise overlaps of TAD boundaries are shown for all samples of this study, after calling boundaries using hicratio (all reads, *d*=0500). Before TAD calling, the Hi-C matrices were either unprocessed (filtered) or corrected using iterative correction (IC) (resolution =40kb).

**Supplementary Figure 7.**
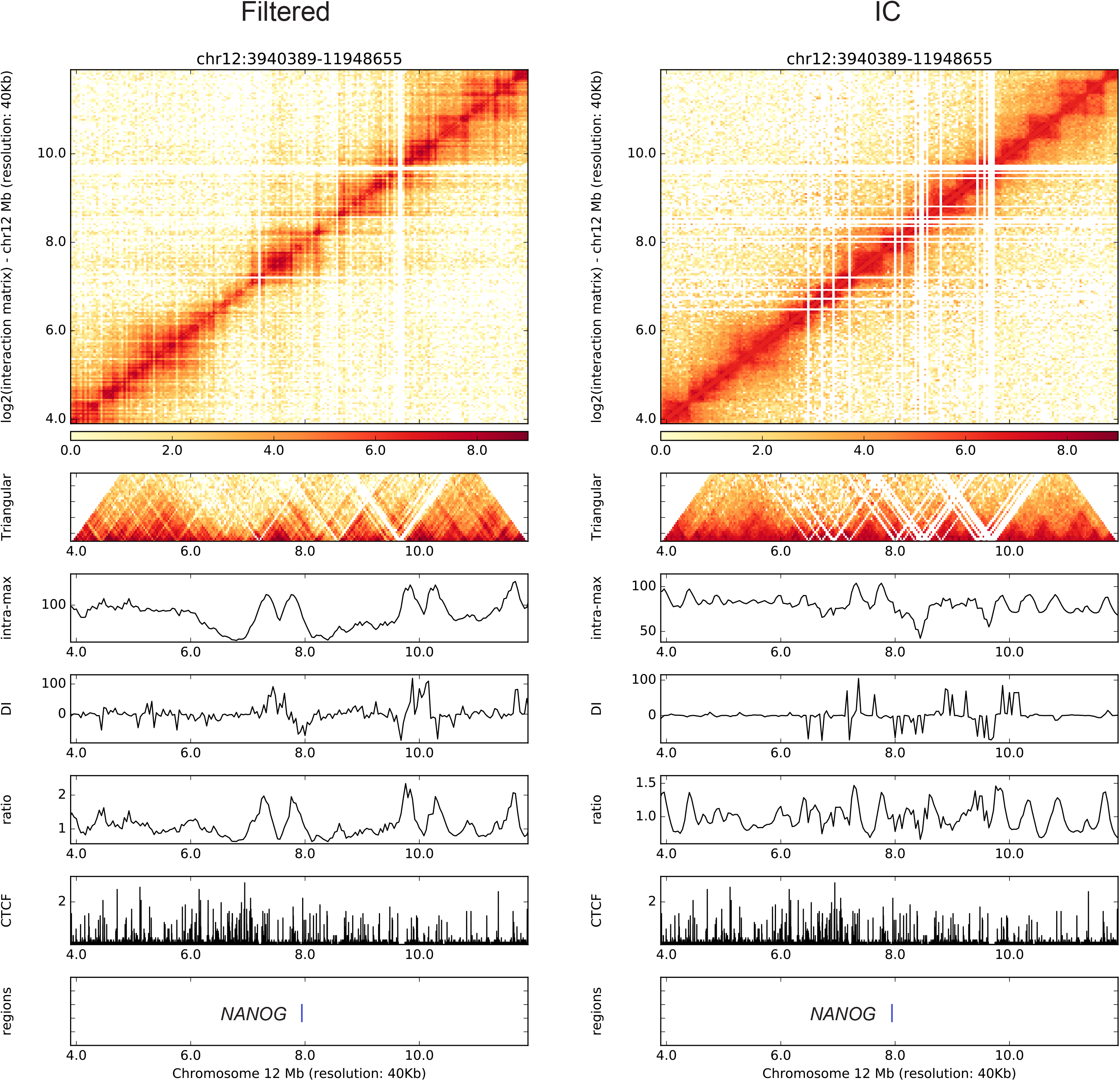
Visualization of TADs and certain areas of interest. HiC-bench integrates HiCPlotter [23] and it offers the ability to easily prepare publication-quality figures. We present the area surrounding *NANOG*, a gene of particular importance for the maintenance of pluripotency. The Hi-C matrix corresponding to the chr12:3940389-11948655 genomic region is presented for H1 cells, before and after matrix correction. The matrix is also rotated 45^o^ to facilitate TAD visualization. Various boundary scores (intra-max, DI, ratio) are shown as individual tracks along with CTCF binding. The location of *NANOG* is presented as a blue line.

**Supplementary Figure 8.**
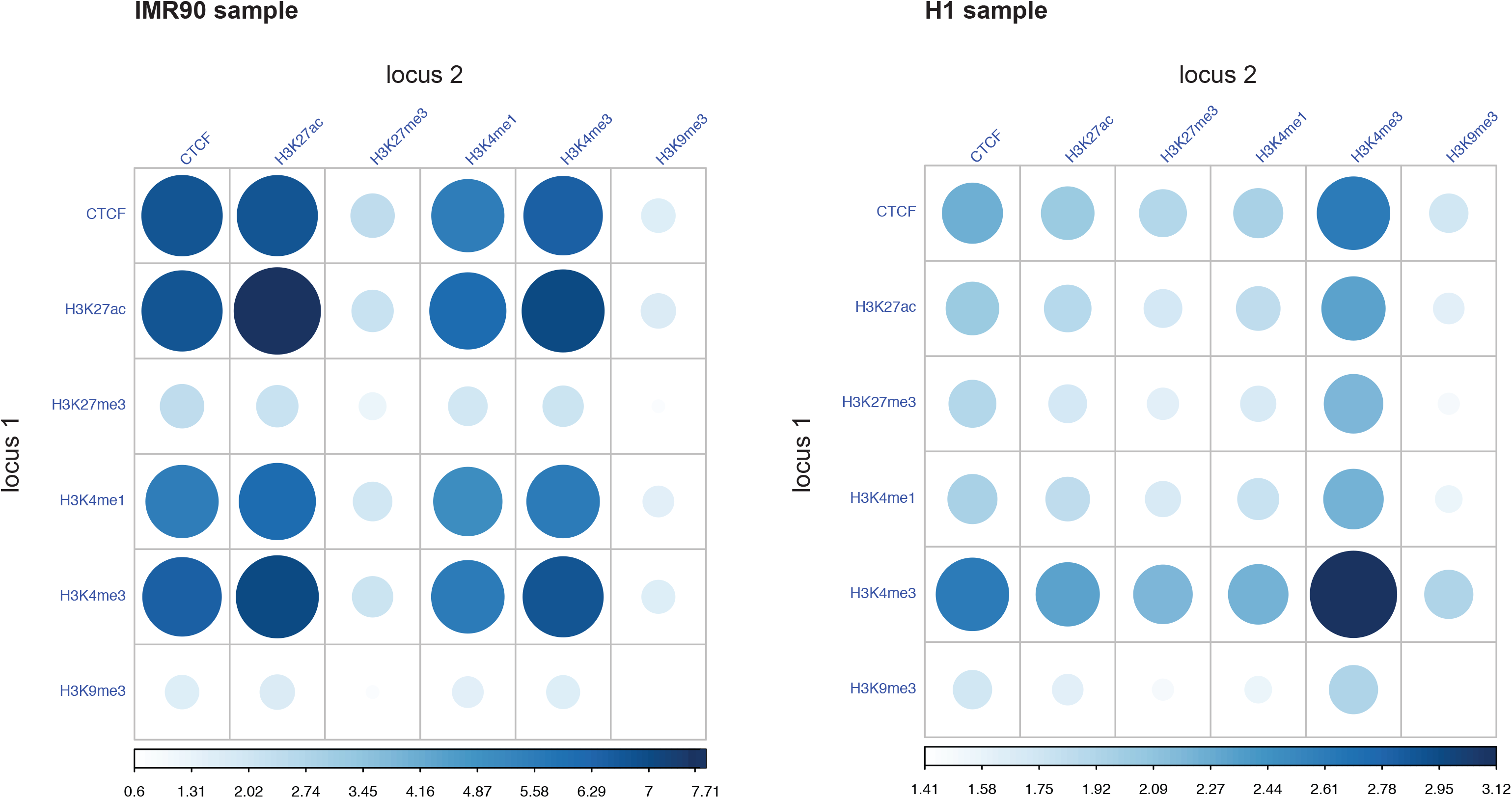
Enrichment of chromatin interactions in human fibroblasts (IMR90) and embryonic stem cells (H1). The enrichment of certain chromatin marks and CTCF in the top 50000 chromatin interactions in the IMR90 and H1 samples is shown. Deep blue and larger circle size indicate higher enrichment.

**Supplementary Table 1. HiC-bench task implementation.** The table summarizes how the pipeline tasks are implemented, which are the requirements for their execution and how they are handled by the *pipeline-master-explorer* script. The first column lists all the tasks performed by pipeline ranging from alignment to annotation. The second column lists the input directory required for each task while the third one lists the parameter files. Certain tasks depend on the reference genome (human or mouse), thus the genome assembly acts as split variable (column 4). In some tasks, replicates can be grouped using the group variable (column 5). Pairwise comparisons between replicates or samples are also allowed using tuples (column 6). The last column lists the full pipeline-master-explorer command for each pipeline task.

**Supplementary Table 2. HiC-bench input-output objects.** The table summarizes the inputs and outputs of the TAD-calling task using three different methods with parameter values stored in the params files (column 2). The first column describes the tree structure of the input directories that are essentially the different Hi-C matrices for each sample, before (filtered) and after matrix correction using different methods (e.g. IC). The second column lists all the different parameter scripts and the third column corresponds to the tree structure of the generated output objects.

